# Fungal endophytes from salt-adapted plants confer salt tolerance and promote growth in Wheat (*Triticum aestivum* L.) at early seedling stage

**DOI:** 10.1101/2021.12.02.470908

**Authors:** Manjunatha Nanjundappa, Nayana Manjunatha, Hua Li, Krishnapillai Sivasithamparam, Michael G K Jones, Ian Edwards, Stephen J Wylie, Ruchi Agarrwal

**Affiliations:** Plant Biotechnology Research Group - Virology, Western Australian State Agricultural Biotechnology Centre, School of Veterinary and Life Sciences, Murdoch University, Perth, Western Australia; Division of Seed Technology, ICAR-Indian Grassland and Fodder Research Institute, Jhansi- 284003, India; ICAR-National Research Centre on Pomegranate, Solapur-413255, Maharashtra, India

**Keywords:** endophytes, salt tolerance, halophytes, growth promoting activity, *Microsphaeropsis arundinis*, wheat

## Abstract

With increasing human global population, increased yield under saline conditions is a desirable trait for major food crops. Use of endophytes, isolated from halophytic hosts, seems to be an exciting approach for conferring salt tolerance to a salt sensitive crop. Therefore, in the current study, fungal endophytes were isolated from halophytic plants’ roots and their ability to withstand in vitro salt stress was evaluated. They could withstand upto 1M NaCl concentrations and this tolerance was independent of their host or tissue source. When inoculated on salt sensitive wheat seeds/seedlings several of the endophytes showed a positive impact on germination and biomass related parameters upon salt stress, both *in vitro* and under glasshouse conditions. One of the isolate from dicot plants (identified as *Microsphaeropsis arundinis*) could successfully colonize wheat and promote its growth under salt and no salt conditions. Amongst the fungal isolates that are known to be natural endophytes of wheat, Chaetomium globosum was the best performing isolate which has been reported as an effective biocontrol agent earlier. Based on the results of our preliminary study, we suggest that these fungal endophytes could prove beneficial for salt stress tolerance enhancement of wheat crop.

---

Soil salinity is considered the scourge for plant growth and crop productivity worldwide^1^. Approximately 1125 m ha of land throughout the world is affected by high levels of salt due to intensive agriculture and desertification processes^2^. Increase in salinity tolerance for the world’s two major crops, wheat and rice, is an important goal as the world’s population is increasing more rapidly than the area of agricultural land^3^. Seed germination and seedling growth of wheat, like other crops, has been found to be negatively affected by salinity stress^4,5^. As a consequence, plant tolerance to salt, mainly to the sodium cation (Na^+^), is a desirable trait to be selected in cultivated crop plants. To overcome salinity stress, tolerant variety can be developed through agronomical and breeding or advanced molecular techniques, but these are time consuming and highly expensive. In this regard, one of the alternative approaches to achieve normal plant growth under salt stress is the efficient utilization of endophytes^6^.

Endophytes (endo = within, phyte = plant) represent an important component of the plant microbiome and comprise of both bacteria and fungi. They are present in all plant species asymptomatically but often promote host performance in terms of growth and resistance to abiotic and biotic stresses. Endophytes isolated from plants growing in warm soils and coastal saline soils indicate a high commercialization potential in agriculture by providing increased crop yield in hot and salty water environments, respectively^7,8^. These previous studies collectively show positive effects of endophytes on improving plant fitness and survival under the different stress conditions, supporting the hypothesis that the effects of endophytes on plant salt stress mitigation may be general among different plant taxa and stress conditions. However, a well-structured study is needed to test this hypothesis.

To draw overall conclusions about the positives of endophytes for plant salt stress tolerance, it is imperative to identify host-endophyte combinations that yield tolerance to salt. In this regard, we isolated the endophytic fungi associated with halophytic plants growing in coastal areas of Western Australia and evaluated their ability to tolerate NaCl stress. The isolates which were tolerant to high concentrations of salt (1 M NaCl), were inoculated on seeds of salt-sensitive wheat germplasm line to examine their ability to confer salt tolerance to the new host. The results of the current study are important because they not only open up exciting possibilities of using endophytes from salt adapted plants for mitigating salt stress in agricultural crops but also in understanding the underlying biochemical and molecular basis of plant-endophyte interaction.

## MATERIALS AND METHODS

### Collection site and sampling

Halophytic plants of eight species were collected from wild populations growing at three coastal sites in Western Australia. Roots and rhizosphere soil of *Oxalis pes-caprae* (soursop, *Oxalidaceae*), *Chenopodium album* (fat-hen, *Amaranthaceace*), *Elymus repens* (couch grass, *Poaceace*), and an unidentified brassicaceous plant (*Brassicaceae*) were collected at Collins Pool, Birchmont, located beside an estuary. Roots and rhizosphere soils of *Salicornia quinqueflora* (beaded samphire, *Amaranthaceace*), *Juncus acutus* (rush, *Juncaceae*), and an unidentified grass (*Poaceace*) were collected at Herron Point, Birchmont, located beside the same estuary. Rhizosphere soil and stolons of *Ammophila arenaria* (marram grass, *Poaceace*), and rhizome of *Posidonia australis* (sea grass, *Posidoniaceae*) were collected from a beach located near the city of Bunbury, Australia (Table S1).

### Measurement of soil salinity and pH

Soil salinity and pH were calculated in the field (Table S1). Five ‘5 cm diameter’ cores of soil were collected adjacent to sampled plants, with the exception of the seagrass samples. Cores were taken to 10 cm depth and thoroughly mixed. A sample of 20 g of soil was placed in a vessel and 100 mL of distilled water was added. The mixture was shaken periodically over one hour, then allowed to stand for 30 min before measuring the salinity (calculated from electrical conductivity) and pH using an EC8500 portable pH and conductivity meter (Apera Instruments, Ohio) according to the manufacturer’s instructions. A temperature compensation coefficient of 2%/°C was used for calculating salinity. pH measurements were later confirmed in the laboratory using an Orion Star A111 pH meter (Thermo Fisher Scientific, Massachusetts).

### Isolation and culture of fungal endophytes

Plant samples were rinsed under running tap water to remove surface debris and soil particles. Fungal endophytes were isolated using a protocol described before^9^. Petri plates were incubated at 25°C for 48 h and fungal colonies were counted. Colonization frequency was estimated as follows:

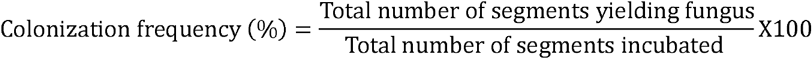

To obtain pure cultures of each fungal isolate, hyphal tips of colonies were transferred to Petri plates containing 0.2x potato dextrose agar (PDA) supplemented with streptomycin sulfate (0.1 mg mL^-1^). Cultures were stored long-term at -80°C in 15% (v/v) glycerol.

### Identification of fungal endophytes

Morphological *viz*., colony color, mycelial texture and growth rate; and margin characteristics were recorded for pure fungal cultures. If several isolates of similar appearance were available from the same host plant then only two were chosen for molecular identification. Genomic DNA was extracted^10^ and Internal Transcribed Spacer (ITS) regions were amplified by PCR using universal primers ITS1 and ITS4 or ITS4 and ITS5^11^. Amplified products were quantified and sequenced. Further, the obtained ITS sequences were compared with those available on databases such as GenBank (NCBI) and UNITE^12^ in order to reveal their identity. Isolates were identified to the species level if their ITS sequences shared ≥97% pairwise similarity with a named species from the databases analysed. When the similarity percentage was 95-96%, only the genus name was accepted and for sequence identities <95%, isolates were classified to the level of family (if available) or labelled as ‘unidentified fungus’ as described earlier^13^. Sequences were aligned using ClustalW and percent similarity was obtained using the EMBL-EBI (http://www.ebi.ac.uk/Tools/msa/mafft/) platform. Phylogeny was estimated using the Maximum Likelihood (ML) method within MEGA v6.06 (http://www.megasoftware.net/) after that ‘Find Best DNA Models’ was applied to determine the most appropriate model for construction of respective ML phylogenies. Predicted tree branches were supported with 1000 bootstrap replications.

### Evaluation of endophytic fungal isolates for salt tolerance

Fungal isolates from each plant species were evaluated for tolerance to salt *in vitro*. Endophytic fungal isolates were sub-cultured on PDA and allowed to grow for 7 d. A 5 mm^2^ agar plug of mycelium was excised from the edge of the colony and used to inoculate potato dextrose salt agar (PDSA) plates, which were PDA plates amended with 1.0 M NaCl. Fungal colonies grown on PDA plates served as control. Cultures were incubated at 25°C in the dark. Three replications were maintained for each treatment. The diameter of each mycelial colony was recorded on the seventh day following plate inoculation. Diametrical growths of colonies were measured at three different diameters per plate and the mean of these measures for overall replications was calculated. Inhibition of growth under treatment on PDSA medium was calculated as a percentage of growth of the same isolate growing on PDA medium. Classification of salt tolerance of endophytic fungi was as described previously^14^. Highly tolerant fungal endophytes were used for further studies.

### In vitro screening of fungal endophytes for conferring salt tolerance to host

The wheat genotypes obtained from Edwards’s laboratory, SABC, Murdoch University, Perth, Australia were initially screened for tolerance at different salt concentrations. All wheat genotypes were found highly sensitive to salt (NaCl) at 150 mM concentration (data not shown), therefore this concentration was used in the current study. Among the wheat genotypes GP#15, with agronomical superiority, was used for further studies. The selected fungal endophytes were evaluated *in vitro* for their ability to impart salinity tolerance to a salt-sensitive wheat genotype at 150 mM NaCl. Two different methodologies were employed for the *in vitro* stress tolerance studies: agar media based method and filter paper methods as illustrated in Figure 1.

**Fig. 1.**
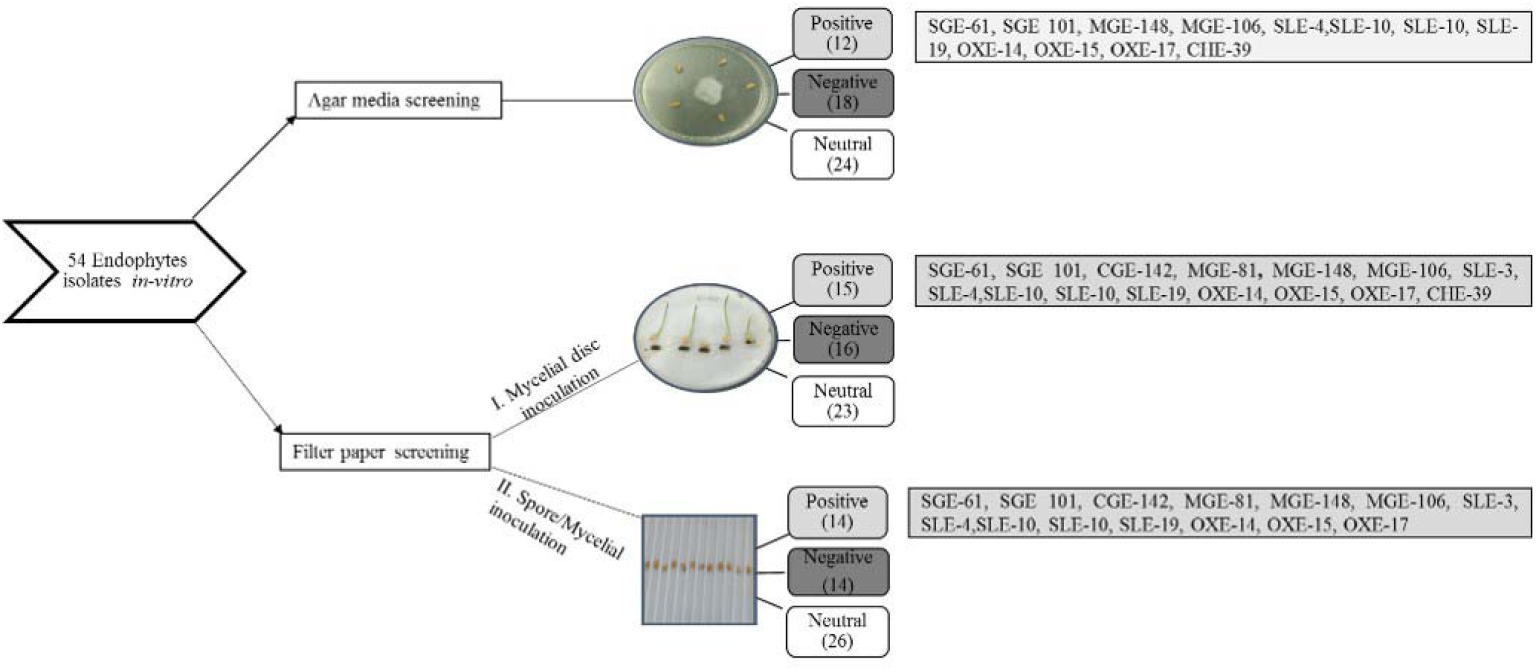
Schematic representation of different approaches employed for *in vitro* screening of fungal endophytes, isolated from halophytic species, that conferred salt (150 mM NaCl) tolerance to wheat seedlings (observation were made at 10 days after treatment). The number in the parenthesis indicates counts of the endophyte isolates showing respective interaction: **Positive**= enhanced tolerance, **Negative** = decreased tolerance or retarded growth upon inoculation, **Neutral** = no measurable influence. Names of the isolates have been provided in the box

#### i. Agar media based test

Each fungal endophyte was applied individually to wheat (GP#15) seeds according to method described^15^. Briefly, a 5 mm^2^ agar plug, cut from the margins of the 10 day old colony were placed, hyphal side down, in the middle of each Petri dish containing 2/4th strength PDA media amended with 150mM NaCl (each Petri dish was filled 2/3rd with media and assumed variation was minimized by replication) and then incubated at 25°C. Five surface sterilized seeds were placed at a distance equivalent to 24 h and 72 h grown culture plates for fast growing and slow growing isolates respectively. The Petri dishes were sealed with parafilm and then incubated at room temperature (25°C). 10 days post-inoculation (dpi), the parafilm was removed and observations on host endophyte reaction were recorded. The control treatments contained no fungus. Each fungal inoculation or control was replicated at least three times.

#### ii. Filter paper□based test

Long strips were made from filter paper (Whatman 10312209, Grade 598) and two strips per furrow were placed in plastic plate with 12 furrows as shown in Figure 1. Sodium chloride solution (250 µl of 150 mM NaCl) was added in each furrow and surface sterilized wheat seed were placed in center of each furrow. The plates were sealed with parafilm (Parafilm^®^ M, P7793, Sigma) and incubated at 25°C in an inclined position to facilitate downward root movement. Once radicle had grown 5 mm in length, a 3 mm^2^ agar plug made from the growing edge of endophytic fungal colony was placed along the radicle of each seedling and plates were sealed again with parafilm and incubated as before. At 10 dpi, the parafilm was removed and host-endophyte reaction was recorded. The control treatments with an agar plug contained no fungus.

Using agar media based method, germination kinetics parameters such as germination percentage (G%) and mean germination time (MGT) and biomass related parameters such as root length, shoot length and seedling fresh weight were recorded for hosts inoculated with endophytes (n=13), referred to as endophyte inoculated (EI) seeds hereafter. Control plates contained non-inoculated (NI) wheat seeds placed on media with or without salt.

### Root colonization by fungal endophytes

Trypan blue (0.01% w/v) staining was used to identify fungal mycelium within root tissues using a method used earlier^16^ with suitable modifications applicable to root tissues. Briefly, seedlings inoculated on filter paper (Method I and II) were collected 15 dpi. Roots were cut into approximately 0.5 cm segments and were cleared with acetic acid:ethanol (1:3 v/v) solution for 12 h. A second tissue clearing was done by soaking tissues in acetic acid:ethanol:glycerol (1:5:1 v/v/v) solution for 5 h. The samples were subsequently incubated overnight in a staining solution of trypan blue. Stained tissues were rinsed with 60% sterile glycerol and stored in it until examination. Specimens were examined under an Olympus BX 51 optical microscope (Olympus, Japan). Five to ten segments were assessed per endophyte inoculation treatment.

### Glasshouse based evaluation of fungal endophytes for conferring salt tolerance to host

Based on the ability of isolates in conferring salt tolerance to wheat *in vitro*, 11 isolates were further selected (SLE-6, SLE-10, SLE-19, OXE-14, OXE-17, SGE-60, SGE-61, MGE-81, MGE-106, MGE-148 and CGE-142) for glasshouse experiment. Spore suspensions were prepared from 10 day old cultures of highly salt tolerant fungal isolates growing in 2/4th strength potato dextrose broth and incubated on a shaker. The mycelial pellicle was washed in sterile water to remove residual broth, then macerated in a blender and filtered through sterile cotton wool. The number of spores was counted using a haemocytometer and diluted to 1×10^7^ spores mL^−1^. Wheat seeds were soaked in spore suspension, of individual endophyte, overnight after which they were taken out from the suspension and shade dried. Five seeds were sown in perforated pots filled with perlite and sand (3:2). The pots were placed in plastic trays either containing 150 mM NaCl solution or water. Each tray contained 6 pots and 500 ml salt solution or water (each pot served as one replication). Similarly seeds soaked in water were sown in six separate pots and placed in a tray containing water and served as control. Once in 3 days salt solution was replaced with fresh salt solution (to avoid salt accumulation in trays they were washed thoroughly and solution was replaced). Results were reconfirmed by repetition of experiment.

### Evaluation of physiological and biomass related parameters of host

#### i. Chlorophyll content (CC)

Chlorophyll content was measured from fully expanded leaves (1^st^ leaf as shown in Figure S1) of seedlings by a hand-held chlorophyll meter (CCM-200 plus, Opti-Sciences Inc., Hudson, NH, USA). Three seedlings per pot were investigated. A total of eighteen seedlings were considered from each treatment and averaged value was taken as CC per seedling. Chlorophyll data measurement was carried out 7, 11 and 15 days after stress was imposed, just before the plants were harvested.

#### ii. Relative Water Content (RWC) and Biomass

Fully expanded leaf of wheat seedling was used to estimate RWC. A total of 10 leaves were harvested randomly from six pots in each treatment. The leaves were placed in polythene bags and transported to the laboratory as quickly as possible in order to minimize water losses due to evaporation and were also weighed immediately to obtain fresh weight (Fw). Then, leaves were soaked in distilled water in test tubes for 24 h at 4 °C in the dark, and turgid weight (Tw) was noted. Subsequently, samples were dried in the oven at 70 °C for 24 h, and dry weight (Dw) was measured. RWC of seedling was determined as:

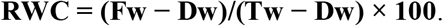

A total of six seedlings were selected per treatment for the estimation of seedling biomass parameters viz., root length, shoot length and root and shoot dry weight. Seedlings were divided into roots and shoots, and soil was washed from roots by hand. Samples were desiccated for 48 h at 80 °C, and dry weight (mg) was recorded. Also, root to shoot ratio was calculated based on their length.

### Statistical analysis

To describe the variability, several simple univariate analysis including means, ranges and variance were calculated. Coefficients of variation (CV%) was also calculated from the variance components and the overall means for all the investigated treatments. Clustering of different treatments based on the CC was carried out using ‘Fastcluster’ package of R statistical software (version 3.4.4) with squared Euclidean distance as a measure of dissimilarity and incremental sums of squares as a grouping strategy^17^. Data of all characters were standardized to a mean of zero and variance of one and Principal Component Analysis (PCA) was performed. First, second and third principal component axes scores were plotted together to visualize the effect of different treatments simultaneously.

## RESULTS

All halophytic plants examined were found to be colonized by multiple culturable fungal endophytes. Two hundred and forty two fungal isolates were obtained from 320 plant specimen and their colonization frequency ranged from 63% to 96% (Figure S2). A high number (96%) of endophytic fungi were isolated from the root tissues of *J. acutus* and *S. quinqueflora* plants (Figure S2). Pure fungal cultures were initially grouped according to their morphological (*viz*., colony color, mycelial texture and growth rate) and margin characteristics.

### Endophytes showed differential response to salinity in vitro

One hundred and thirty fungal isolates were screened for their responses to 1.0 M NaCl *in vitro*. Based on the degree of inhibition of radial growth on PDSA medium compared to PDA medium, fungal isolates were grouped as highly-tolerant, tolerant, moderately-tolerant, or sensitive (Table S2) as described earlier^14^. Most isolates (58) were grouped into the moderately-tolerant category, followed by 39 isolates that were tolerant and 27 that were highly-tolerant. The growth of 6 isolates was severely inhibited on PDSA therefore they were categorised as sensitive. Endophytes originating from monocotyledonous and dicotyledonous halophytes differed in their salt tolerance as shown in the Figure 2. Most isolates from dicotyledonous halophytic hosts had moderate to high salt tolerance whereas most endophytes from monocotyledonous hosts had moderate tolerance. Among the halophytes, sea grass, a species constantly immersed in seawater (∼550 mM NaCl), was colonised with the most highly salt-tolerant endophytes. Moreover, isolates inhabiting the same host also showed differential levels of salt tolerance.

**Fig. 2.**
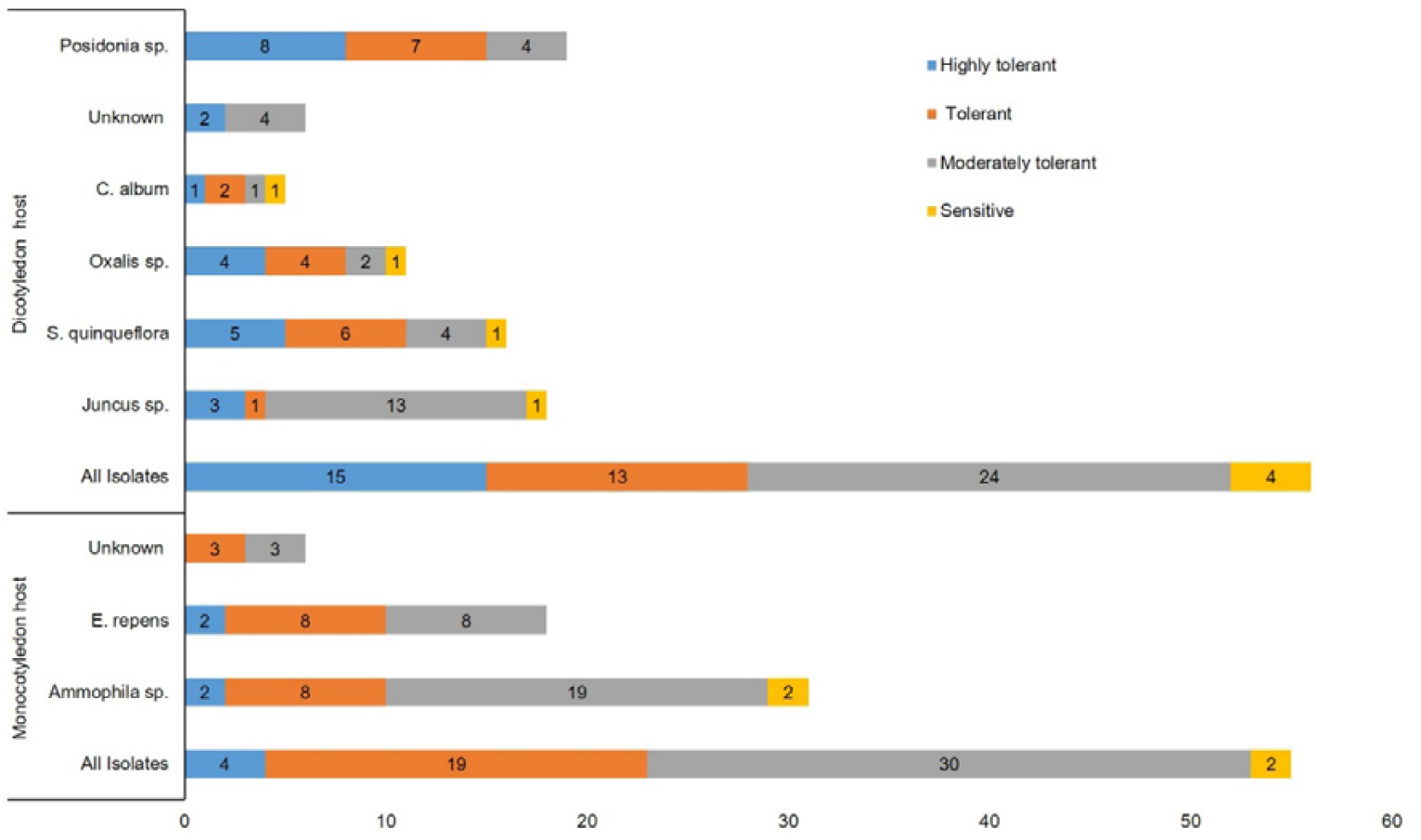
Categorization of endophytes originating from monocotyledonous and dicotyledonous halophytes based on the difference in their salt tolerance

### Fungal endophytes inhabiting halophytes belonged to highly diverse genera

Based on morphological characteristics and inherent salt tolerance, fifty-four representative fungal isolates were selected for molecular identification, where several isolates of similar appearance were isolated from the same host plant, only two isolates were chosen for molecular identification. The chosen isolates were identified based on ITS-amplicon sequencing results followed by database similarity search. The ITS sequences obtained have been deposited in the NCBI GenBank (Accession No. MK431041-MK431094; Table S3). Database similarity search revealed that diverse fungal flora had colonized the halophytic hosts used in the study. Twenty isolates could be identified completely i.e. upto species level with some unidentified to the genus level (Table S3). All endophytes isolated from halophytes were members of phylum Ascomycota and most of them belonged to subphylum *Pezizomycotina* (48), and were distributed in three classes viz., *Dothideomycetes* (17) *Eurotiomycetes* (7) and *Sordariomycetes* (24). Among the fungal orders, *Hypocreales* (15), *Pleosporales* (17) and *Eurotiales* (7) were the most highly represented (Figure 3). Dominant genera identified in this study were *Alternaria, Chaetoium, Fusarium* and *Penicillium*, whereas genera that were represented by only one or a few isolates were *Aquanectria, Aspergillus, Bipolaris, Clonostachys, Didymella, Didymosphaeria, Microascus, Paraconiothyrium, Paraphaeosphaeria, Phaeosphaeria, Phoma, Phomopsis, Plectosphaerella, Setosphaeria, Soradria* and *Trichoderma*. The six isolates that could not be identified were classified as *Incertae sedis*. Further, phylogeny revealed that highly salt-tolerant endophytic isolates grouped into a single cluster (Figure 4) indicating that they may share some similarity at genetic level. No phylogenetic pattern was however evident with regard to plant tissue type or host species (data not shown).

**Fig. 3.**
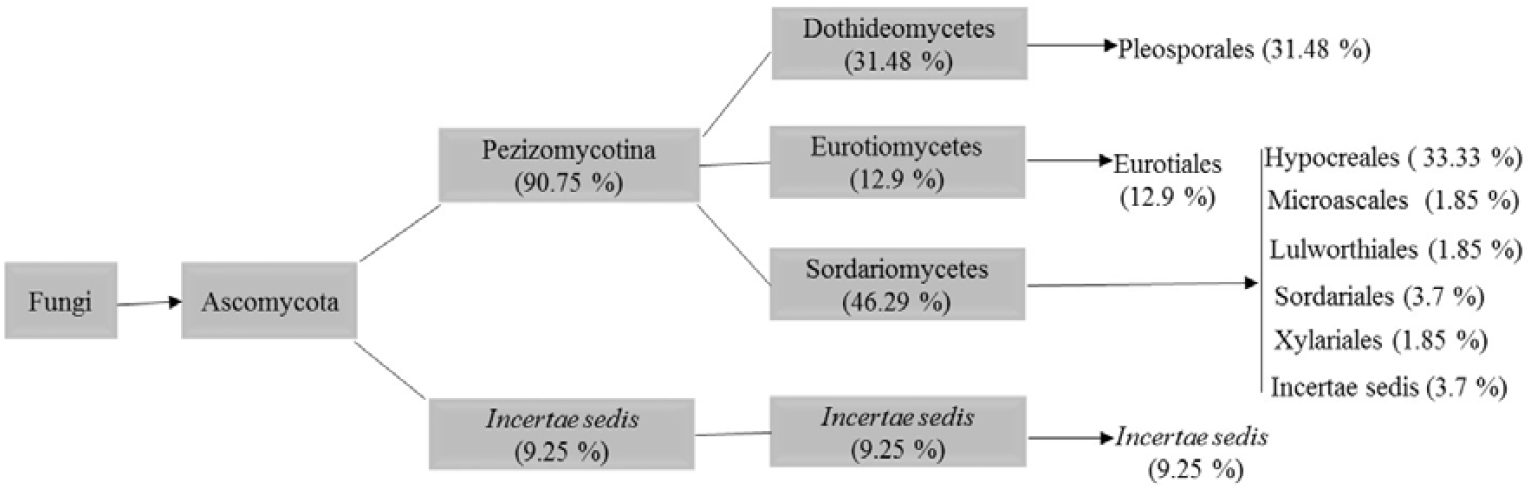
Schematic representation of phylogenetic placement of 54 fungal species identified from ITS sequences of endophytes isolated from different halophytic species. Classification follows Hibbett et al. (2007)

**Fig. 4.**
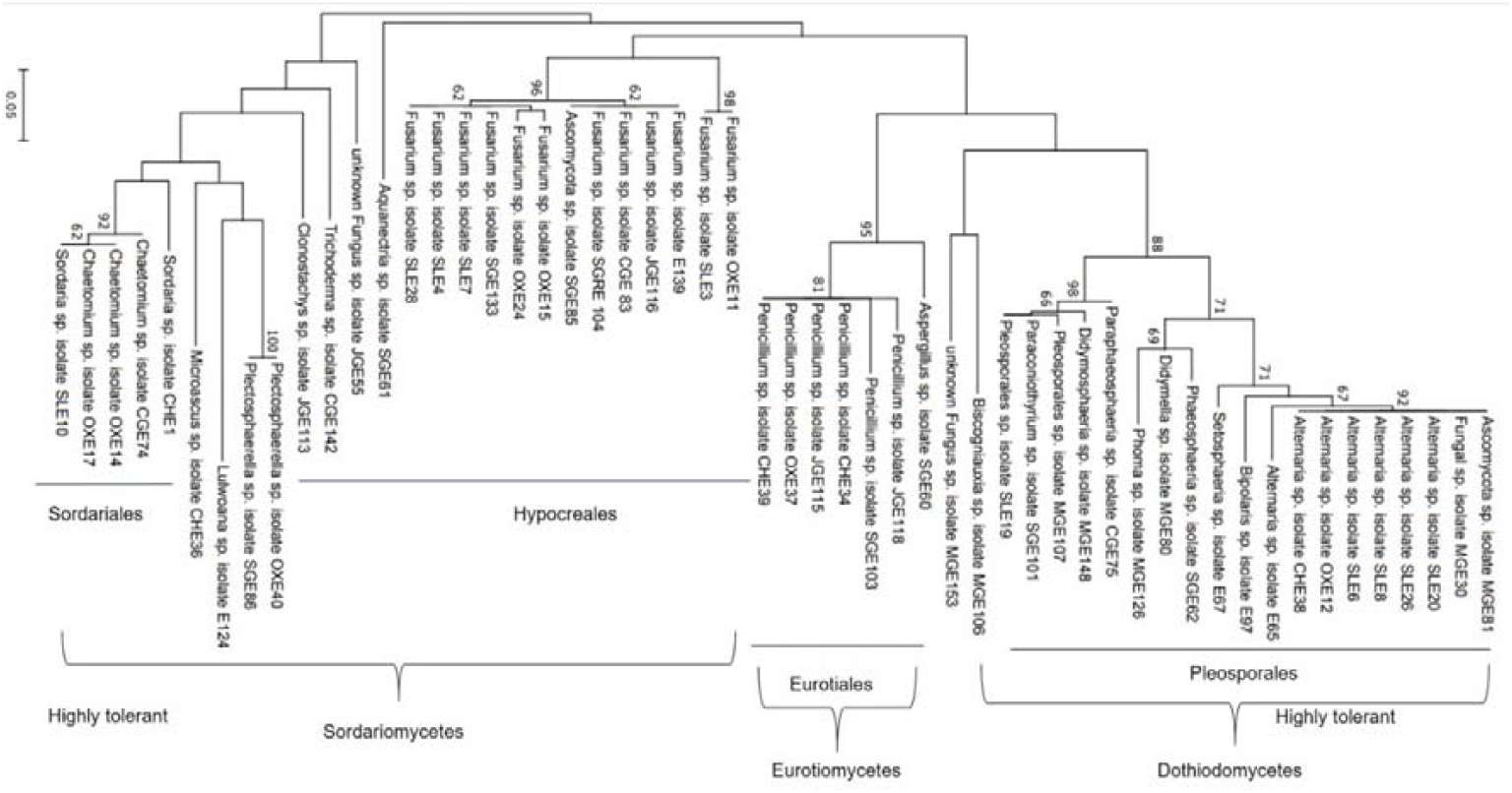
Phylogenetic analysis of fungal endophytes isolated from halophytic hosts revealed that highly salt-tolerant endophytic isolates grouped into a single cluster

### Fungal endophytes isolated from different tissues showed positive impact on salinity tolerance of wheat in-vitro

Seeds of the salt-sensitive wheat genotype (GP#15) were inoculated with 54 endophytes and the performance was evaluated on 150 mM NaCl using different approaches (Figure 1). Isolates enhancing seedling salt stress tolerance had been isolated from all type of tissues used under study. However, on agar method 21 % of isolates from the roots had positive impact whereas on filter paper method, 27 % of isolates from stolons had more positive impact (Figure S3). Based on their positive impact on seedling performance under salinity, some of these isolates were used for further analysis. Roots of EI seedlings were examined under a light microscope for the proof of endophytic colonization. Stained roots highlighted the presence of a network of hyphae, most of which penetrated the intercellular spaces of the root (Figure 5). The pure culture of some of these isolates are shown in Figure S4.

**Fig. 5.**
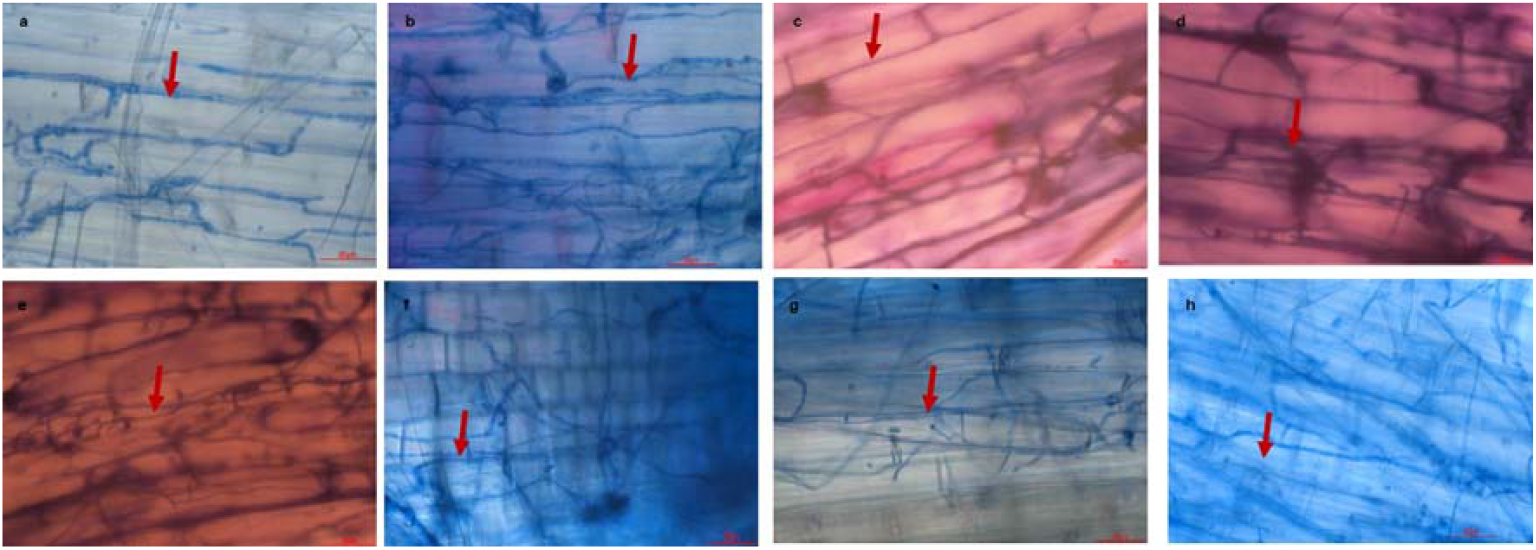
Colonization by the fungal isolates inside the root tissue of salt sensitive wheat inoculated in the filter paper screening test. The red-coloured arrow head indicates the presence of fungal mycelia as observed under a compound microscope stained after trypan blue staining a) *Trichoderma atroviride* b) *Alternaria infectoria* c) *Alternaria chlamydospora* d) *Microsphaeropsis arundinis* e) *Didymosphaeria variabile* f) *Chaetomium globosum* g) *Chaetomium globosum* h) *Chaetomium globosum*

### Fungal endophytes promotes growth of wheat at early seedling stage under saline conditions

Seeds of the salt-sensitive wheat genotype (GP#15) were treated with promising endophytes (n=13) and the performance of the seedlings was analyzed both *in vitro* and under glasshouse conditions after subjecting them to salt stress (150 mM NaCl).

Observations of *in vitro* assay revealed that G% of EI seeds placed on salt containing media (SCM) ranged from 88.8 to 97.7%. The NI seeds showed 91% germination on SCM and 97.7% on media without salt (MWS) indicating a higher germination rate in EI seeds than NI seeds on SCM. The EI seeds placed on SCM showed mean germination time (MGT) lesser than NI seeds on the same media, indicating that EI seeds germinated faster than NI seeds when placed on SCM. However, the least MGT was recorded for NI seeds placed on MWS (Figure 6).

**Fig. 6.**
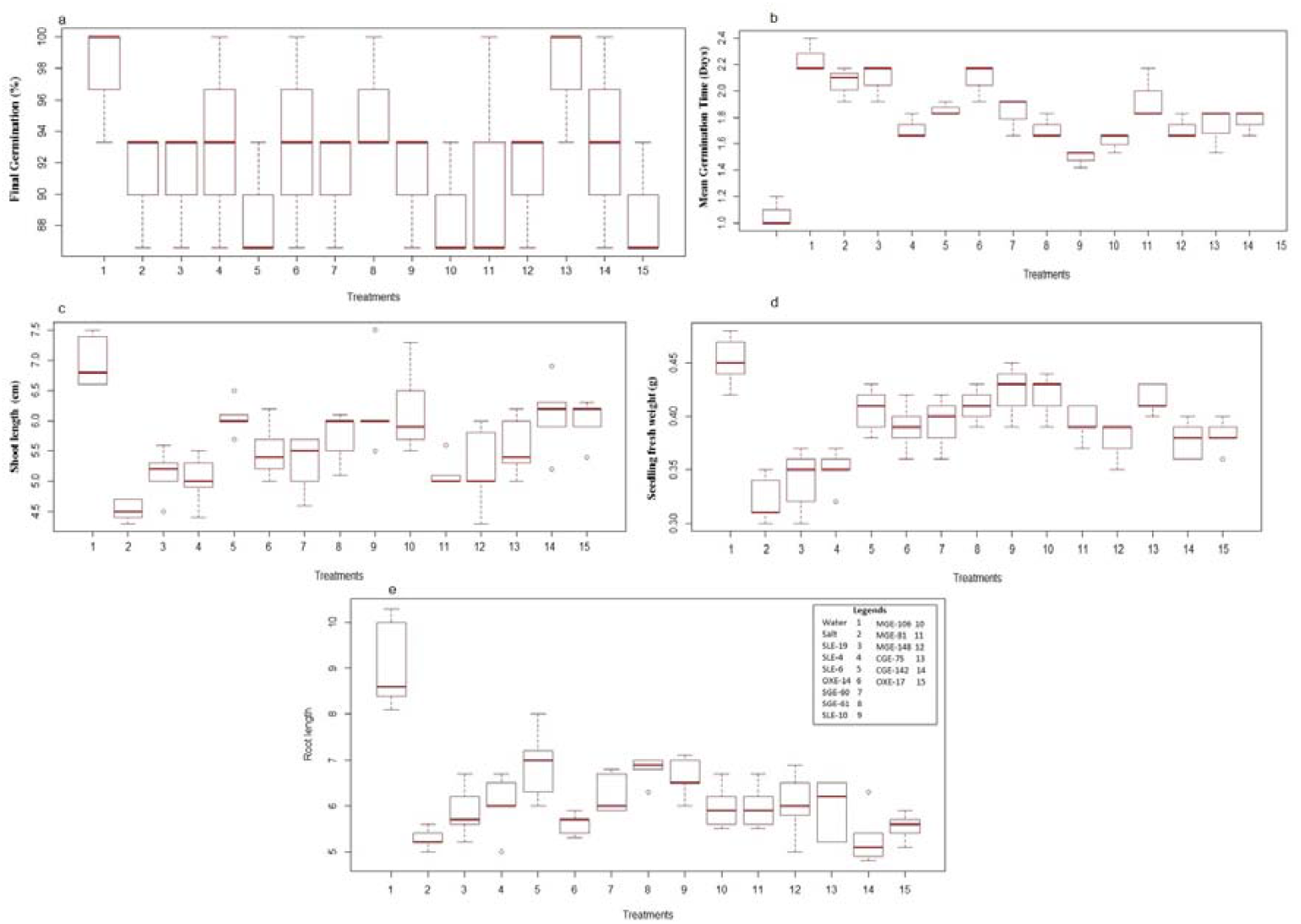
Measurement of germination kinetics parameters and seedling biomass related parameters after subjecting the endophyte inoculated wheat (GP#15) seeds to salt stress (NaCl 150 mM) *in vitro*

Similar trends were observed for seedling biomass related parameters such as root length, shoot length and seedling fresh weight. The EI seedlings showed higher biomass as compared to NI seedlings on SCM (Figure 6). All the endophytic isolates improved the performance of the wheat seedlings under salt stress, however, we selected 11 best performing isolates for the glass house based studies (Table 1).

**Table 1.** Endophytic fungal isolates identified and used in the current study for evaluating their ability to confer salt (150mM NaCl) tolerance to wheat seeds/seedlings *in vitro* or in glass house respectively.

| S. No. | Isolate code | Identity based on ITS sequencing and database similarity search | Host species | Salt tolerance | Accession no. |
| --- | --- | --- | --- | --- | --- |
| 1. | SLE-6 | *Alternaria chlamydospora* | *Salicornia quinqueflora* | Highly tolerant | MK431069 |
| 1. | SLE-19 | *Microsphaeropsis arundinis* |  | Highly tolerant | MK431073 |
| 2. | SLE-10 | *Chaetomium globosum* |  | Highly tolerant | MK431072 |
| 3. | OXE-14 | *Chaetomium globosum* | *Oxalis pes-caprae* | Tolerant | MK431079 |
| 4. | OXE-17 | *Chaetomium globosum* |  |  | MK431081 |
| 5. | SGE-60 | *Aspergillus ochraceus* | *Posidonia australis* | Highly tolerant | MK431046 |
| 6. | SGE-61 | *Aquanectria penicillioides* |  | Highly tolerant | MK431047 |
| 7. | MGE-81 | *Alternaria infectoria* | *Ammophila arenaria* | Highly tolerant | MK431061 |
| 8. | MGE-106 | Unknown fungal sp. |  | Highly tolerant | MK431062 |
| 9. | MGE-148 | *Didymosphaeria variabile* |  | Tolerant | MK431065 |
| 10. | CGE-142 | *Trichoderma atroviride* | *Elymus repens* | Tolerant | MK431058 |

For EI seedlings grown under glasshouse conditions, CC was measured at 7, 11 and 15 days after stress (das) treatment (Figure 7) to study the effect of duration of stress on CC of leaves. Chlorophyll estimation was also performed for NI seedlings grown in solution with or without salt at the same time points. Higher CC was observed in seeds treated with fungal isolate CGE-142, especially as the duration of stress increased (11 and 15 das). Similarly, CGE-142 inoculation enhanced the CC of seeds grown in solution without salt (SWS). These values were higher than the CC of NI seeds grown in SWS or salt containing solution (SCS), indicating that the fungal isolate induced higher chlorophyll synthesis in wheat seedlings.

**Fig. 7.**
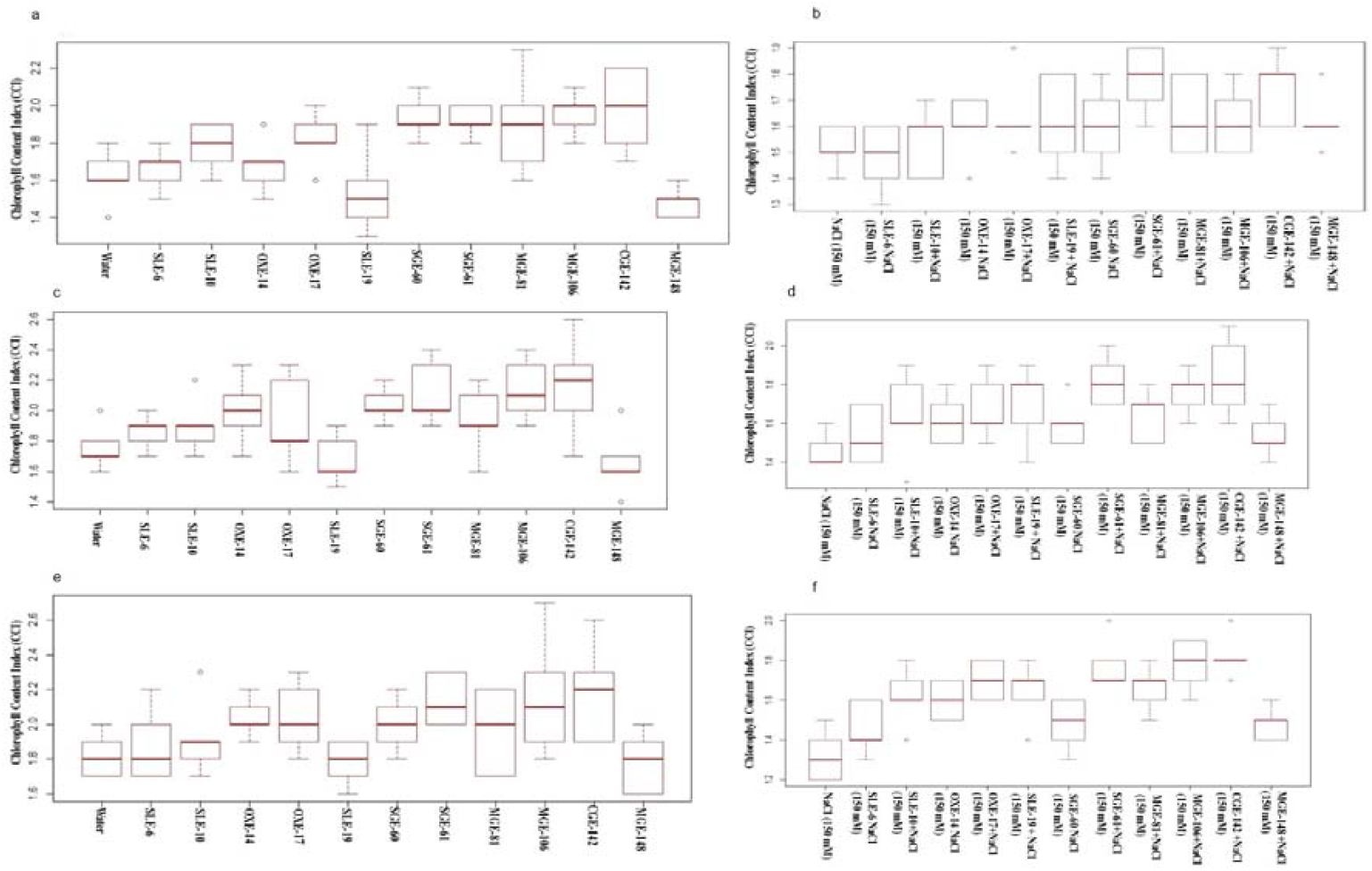
Measurement of chlorophyll content index of endophyte inoculated wheat (GP#15) seedlings at 7 (a, b), 11 (c, d) and 15 (e, f) days of salt stress (b, d, f) or no stress (a, c, e) under glass house condition

All the fungal endophytic isolates improved the RWC of wheat seedlings as compared to NI seedlings when grown in SCS (Table 2). The highest RWC was observed in seedlings inoculated with CGE-142. With regard to biomass-related parameters, seedlings inoculated with SLE-10 exhibited root and shoot biomass (fresh weight, dry weight and length) as well as root to shoot ratio higher than NI seedlings grown in SCS. However, similar trend was observed in seedlings inoculated with CGE-142 in the absence of salt (Table 3).

**Table 2.** Measurement of biomass related parameters of endophyte inoculated wheat (GP#15) seedlings grown under salt stress condition in the glass house

| Isolate code | RWC (%) | % change over control | RDW (g) | % change over control | SDW (g) | % change over control | RL (cm) | % change over control | SL (cm) | % change over control | RSR | % change over control |
| --- | --- | --- | --- | --- | --- | --- | --- | --- | --- | --- | --- | --- |
| SLE-6+NaCl | 95.0ab | 5.62 | 0.07d | -37.15 | 0.08h | -32.88 | 8.76def | 6.07 | 12.4f | -7.83 | 0.74ab | 20.04 |
| SLE-10+NaCl | 93.78b | 4.28 | 0.11a | 0.76 | 0.13cd | 9.96 | 9.70bcdef | 18.69 | 14.8de | 9.87 | 0.66abcd | 10.46 |
| OXE-14+NaCl | 95.0ab | 5.61 | 0.07d | -37.43 | 0.11ef | -5.26 | 9.20cdef | 11.63 | 16.4cd | 21.20 | 0.56def | -8.17 |
| OXE-17+NaCl | 90.94c | 1.15 | 0.08cd | -28.55 | 0.12de | 0.38 | 12.20a | 46.44 | 17.6bc | 30.86 | 0.71abc | 15.27 |
| SLE-19+NaCl | 90.90c | 1.10 | 0.09bc | -19.07 | 0.14bc | 20.27 | 10.30bcd | 23.95 | 18.2bc | 34.56 | 0.57cde | -6.31 |
| SGE-60+NaCl | 95.0ab | 5.63 | 0.08cd | -27.76 | 0.09gh | -23.24 | 8.90cdef | 7.94 | 14.8ed | 8.41 | 0.62bcd | 0.23 |
| SGE-61+NaCl | 90.9c | 1.10 | 0.10ab | -10.84 | 0.10fg | -15.79 | 10.56abc | 27.48 | 17.3bc | 28.43 | 0.62bcd | 1.58 |
| MGE-81+NaCl | 95.82ab | 6.57 | 0.09bc | -18.61 | 0.10fg | -13.79 | 11.10ab | 32.61 | 14.7def | 9.03 | 0.76a | 24.96 |
| MGE-106+NaCl | 96.32a | 7.11 | 0.04e | -63.45 | 0.15ab | 28.96 | 8.06f | -3.77 | 18.9ab | 39.50 | 0.43f | -31.18 |
| CGE-142+NaCl | 96.02a | 6.78 | 0.04e | -63.72 | 0.16a | 35.65 | 9.80bcde | 19.05 | 20.9a | 54.10 | 0.47ef | -22.87 |
| MGE-148+NaCl | 91.28c | 1.48 | 0.07d | -37.43 | 0.10fg | -15.79 | 8.20ef | 0.10 | 13.0ef | -3.97 | 0.63abcd | 3.61 |
| NaCl (150 mM) | 90.0c |  | 0.11a |  | 0.12de |  | 8.36ef |  | 13.6ef |  | 0.62bcd |  |

RWC: Relative water content, RDW: Root dry weight, SDW: Shoot dry weight, RL: Root length, SL: Shoot length, RSR: root to shoot ratio based on length. Same letters indicate statistically insignificant differences ( $p>0.05$ )

**Table 3.** Measurement of biomass related parameters of endophyte inoculated wheat (GP#15) seedlings grown under no stress condition in the glass house

| Isolate code | RWC (%) | % change over control | RDW (g) | % change over control | SDW (g) | % change over control | RL (cm) | % change over control | SL (cm) | % change over control | RSR | % change over control |
| --- | --- | --- | --- | --- | --- | --- | --- | --- | --- | --- | --- | --- |
| SLE-6 | 88.29b | 11.31 | 0.12bc | 7.09 | 0.13de | -12.58 | 12.6e | **-9.98** | 14.8de | -6.29 | 0.85cde | -3.48 |
| SLE-10 | 88.39b | 11.44 | 0.12bc | 11.09 | 0.14cd | -6.24 | 14.9dc | 6.21 | 15.0de | -4.92 | 1.01bc | 13.78 |
| OXE-14 | 88.57b | 11.67 | 0.14a | **29.54** | 0.13e | -13.89 | 16.3bc | 16.61 | 14.3de | -9.28 | 1.18a | **32.37** |
| OXE-17 | 88.37b | 11.42 | 0.13ab | 17.43 | 0.16ab | 5.52 | 13.58de | -2.78 | 18.2bc | 15.30 | 0.75de | -15.36 |
| SLE-19 | 91.89a | 15.88 | 0.12bc | 7.09 | 0.12e | **-21.20** | 17.2ab | 22.82 | 16.0cd | 1.18 | 1.07ab | 20.91 |
| SGE-60 | 92.31a | **16.42** | 0.13ab | 17.43 | 0.13de | -12.83 | 14.9dc | 6.00 | 16.8bcd | 6.14 | 0.91c | 3.04 |
| SGE-61 | 88.37b | 11.43 | 0.14a | 27.49 | 0.14cd | -6.24 | 17.3ab | 23.64 | 18.8b | 19.13 | 0.94bc | 5.58 |
| MGE-81 | 84.61c | 6.69 | 0.14a | 27.09 | 0.16ab | 5.52 | 17.4ab | 23.76 | 18.06bc | 14.32 | 0.96bc | 8.26 |
| MGE-106 | 88.56b | 11.67 | 0.12bc | 6.43 | 0.17a | **13.37** | 13.4de | -4.15 | 18.8b | 19.09 | 0.71e | **-19.30** |
| CGE-142 | 91.85a | 15.83 | 0.13ab | 16.98 | 0.16ab | 5.52 | 18.6a | **32.80** | 21.9a | **38.91** | 0.85cde | -3.97 |
| MGE-148 | 91.67a | 15.59 | 0.11c | 0.22 | 0.12e | **-21.20** | 12.64e | -9.73 | 13.0e | **-17.72** | 0.99bc | 10.76 |
| Mock | 79.31d |  | 0.11c |  | 0.15bc |  | 14.04de |  | 15.8cd |  | 0.89cd |  |

RWC: Relative water content, RDW: Root dry weight, SDW: Shoot dry weight, RL: Root length, SL: Shoot length, RSR: root to shoot ratio based on length. Same letters indicate statistically insignificant differences (p>0.05)

## DISCUSSION

The ability to tolerate briny water is essential for wild plants that live in coastal and marine environments. Such halophytic plants appear to associate widely with fungi as evident from the wide range of fungi described from marine-influenced systems of coastal sand dunes, mangroves, seagrass and estuaries. Although, the roles played by these fungal endophytes in salt tolerance of halophytes could be dependent on several factors^18^ it can be expected that their association with such hosts would provide them salt tolerance too^19^.

Therefore, in the current study, we isolated fungal endophytes from halophytic plants and tested their response to a high-salt environment *in vitro*. We challenged the endophytes with a NaCl concentration (upto 1M NaCl) almost twice that of seawater. This concentration reduced the growth rate of all isolates, but for many, growth inhibition was <50% that of low salt conditions, an indication of tolerance to high osmotic gradients. We are aware that the environment on PDSA medium is not likely to be equivalent to conditions in the interstitial spaces between cells of salt-tolerant plants due to two reasons. Firstly, on a solid medium mycelium is not immersed in an aqueous environment. Secondly, plants actively pump Na^+^ and Cl^-^ ions from the roots, they compartmentalize salts in vacuoles, and they develop osmotic tolerance (involving long-distance signalling) to cope with saline environments^20^. Thus, the salinity experienced by the fungus within the plant may be less than the external salt concentration. Hence, if the fungal isolates could tolerate high salt concentration *in vitro*, they can be expected to tolerate salt stress *in planta* also. As reported earlier, the levels of tolerance to salt by endophytes mainly depends on genetic factors of the fungus species and host habitat^21^, the range of external salinity and accumulation of osmo-protectants in the cytoplasm^18,22,23^. Therefore, the observed variations in salt tolerance of fungal endophytes isolated from different hosts as well as the same host could be due to their genetic constituents or their interaction with halophytic hosts. This finding is in line with earlier studies that showed halophyte microbiomes exhibit differential levels of salt tolerance^24,25,26^. Moreover, all the highly salt tolerant isolates grouped together in a single cluster upon phylogenetic analysis indicating underlying similarities between them.

It is unclear if salt tolerance of fungal endophytes isolated in the current study corresponded to their possible roles in mediating salinity tolerance in their naturally salt-adapted host plants, but it would have been interesting to observe their effect on growth of salt-sensitive crop plants. Therefore, we selected 54 isolates, based on their inherent salt tolerance, and inoculated them on salt sensitive wheat (GP#15) seeds. Mostly they exerted a highly positive impact on GP#15 as evaluated by various *in vitro* techniques. These isolates were then identified based on their ITS region sequences and found to belong to diverse genera (Figure 3, Table S3). Earlier, endophytes isolated from mangrove leaves have been reported to belong to *Acremonium, Phomopsis, Phyllosticta*, and *Sporormiella*^27^, *Diaporthe*^28^, *Bruguiera*^29^, *Aspergillus* species and others^30^. Only a few of these genera were represented in our study, the difference could have been due to different tissue from where endophytes have been isolated, for example we isolated endophytes from roots, however the earlier report had used leaves of halophytic plants.

Further, some of these isolates (n=11) were selected for evaluating their growth promoting activity on GP#15 when subjected to salt stress *in vitro* and under glasshouse conditions. Their effect on germination kinetics of GP#15 seeds *in vitro* was also recorded. Germination related parameters such as G% and MGT were better for EI wheat seeds than NI seeds placed on SCM. Our results also showed that the endophytes inhabiting wheat plants not only promote the growth of the seedling but also confer salt tolerance to the host as compared to the NI control. This was found to be associated with an altered CC and RWC of the wheat seedlings as well as enhanced biomass. Notably, the endophyte treatment provided an advantage to the wheat seeds with regard to their germination and biomass related parameters when exposed to saline conditions. Endophytic association is a promising approach to enhance salt tolerance, although specific mechanisms for this are unclear. The presence of endophytes may stimulate inherent plant responses to salinity and/or provide fungus derived compounds that mediate the stress response. In barley, the presence of the endophyte *Piriformospora indica* induced elevation of ascorbic acid and antioxidant enzymes in roots under salt stress^31^. *Phoma glomerata* and *Penicillium* sp. endophytes in dwarf rice secreted gibberellic acid and indole acetic acid to promote growth under saline conditions^32^.

The basis for selecting these 11 isolates was mainly driven by their salt tolerance ability, however we also tried to include isolates that have never been reported to inhabit wheat naturally. Interestingly, when the roots of EI seedlings were observed under microscope they were found to be colonised by these fungal isolates successfully. Therefore, our study reports for the first time, the colonization of wheat roots by *Microsphaeropsis arundinis, Aspergillus ochraceus, Aquanectria penicillioides, Didymosphaeria variabile*. Among these isolates the best growth promoting activities were exhibited by *Microsphaeropsis arundinis* under both salt and no salt conditions in the glasshouse. This endophyte was isolated from a dicot host but when inoculated on wheat (a monocot), it could not only colonize the new host successfully (Figure 5d) but also promote its growth irrespective of the presence of salt in the growing medium. Amongst the fungal isolates that are known to be natural endophytes of wheat, *Chaetomium globosum* was the best performing isolate. This fungus has been reported to be useful as a bio-control agent against a broad range of pathogens or insect pests^33, 34^. Moreover, its effect on wheat seedlings under drought conditions has also been reported^35^. Therefore, in addition to these reports, based on the results of the current study we suggest that *Chaetomium globosum* may be useful in conferring resistance to biotic stress as well as tolerance to abiotic stress to wheat seedling. The current work represents the first step in the process of identifying candidate endophytes to partner with agricultural plants threatened by saline soils. However, the biochemical and molecular mechanism for conferring salt tolerance to new host needs to be elucidated in future. Furthermore, future studies should be focused on application of potential salt tolerant isolates obtained in this study either alone or in combination to impart salt tolerance in different cereal crops under varied salt concentrations.

## CONCLUSION

The amount of salt-affected agricultural land is expected to increase globally in response to climate change. Progress towards increasing crop tolerance to salt through traditional breeding has had limited success, largely because of the genetic complexity of the trait. Endophytes surviving at extreme environmental conditions (dryness, salinity, temperature, heavy metals, etc.) have been found suitable for use in different agricultural practices to combat the effects of such abiotic stress on crop productivity. In the present work, culturable endophytic fungi with different taxonomic affinities were isolated from halophytic plant species. In-vitro studies showed majority of the isolates were found tolerant to high concentration of salt (1.0 M NaCl). When inoculated on seeds of a salt-sensitive wheat genotype (GP#15) these isolates were found to improve germination kinetics of seeds and promote the growth of seedlings under saline conditions.

## Acknowledgements

NM acknowledges the Department of Education and Training, Australian Government for providing a fellowship under the Endeavour Research Program, and Murdoch University for hosting the fellowship. NM and MN thank the Indian Council for Agriculture Research and the Indian Grassland and Fodder Research Institute for granting permission to avail the fellowship program.

