## Supplementary data for "Fungal endophytes from salt-adapted plants confer salt tolerance and promote growth in Wheat (*Triticum aestivum* L.) at early seedling stage"

**Table S1**. Halophytic plants and emergence of endophytic fungi from the tissues of halophytic plants

| Collection site and identies of the halophytic plants (where known) | | | | | | | | Colonization details of endophytic fungi | | | |
|---|---|---|---|---|---|---|---|---|---|---|---|
| Collection site | Plant species collected (if known) | Plant type | Co-ordinates | Soil pH[c] | Soil salinity[c] (ppt) | Water pH | Water salinity (ppt) | Tissue origin | No. of fragments | No. of fungal (isolates) | Colonization frequency (%) |
| Collins Pool, Birchmont | *Oxalis pescaprae* | Dicot | 32º44’30.0” S, 115º42’38.2” E | 8.0 | 40.7 | 8.7 | 31.7 | Root | 28 | 22 | 79 |
| | *Chenopodium album* | Dicot | | | | | | Root | 21 | 15 | 71 |
| | Unidentified brassica | Dicot | | | | | | Root | 25 | 17 | 68 |
| | *Elymus repens* | Monocot | | | | | | Root | 44 | 28 | 64 |
| Herron Point, Birchmont | *Salicornia quinqueflora* | Dicot | 32º42’44.7” S, 115º42’09.1” E | 8.1 | 36.8 | 8.5 | 32.2 | Root | 25 | 24 | 96 |
| | Unidentified grass | Monocot | | | | | | Stolon | 25 | 19 | 76 |
| | *Juncus acutus* | Dicot | | | | | | Root | 28 | 27 | 96 |
| Ocean Drive, Bunbury | *Ammophila arenaria*[a] | Monocot | 33º20’55.9” S, 115º37’14.0” E | 8.3[a] | 24.9[a] | 8.0 | 34.9 | Stolon | 76 | 52 | 68 |
| | *Posidonia australis*[b] | Dicot | | | | | | Rhizome | 48 | 38 | 79 |

[a] soil associated with the *A. arenaria* plant tested.
[b] Fresh plant washed up on the beach
[c] measured in the current study

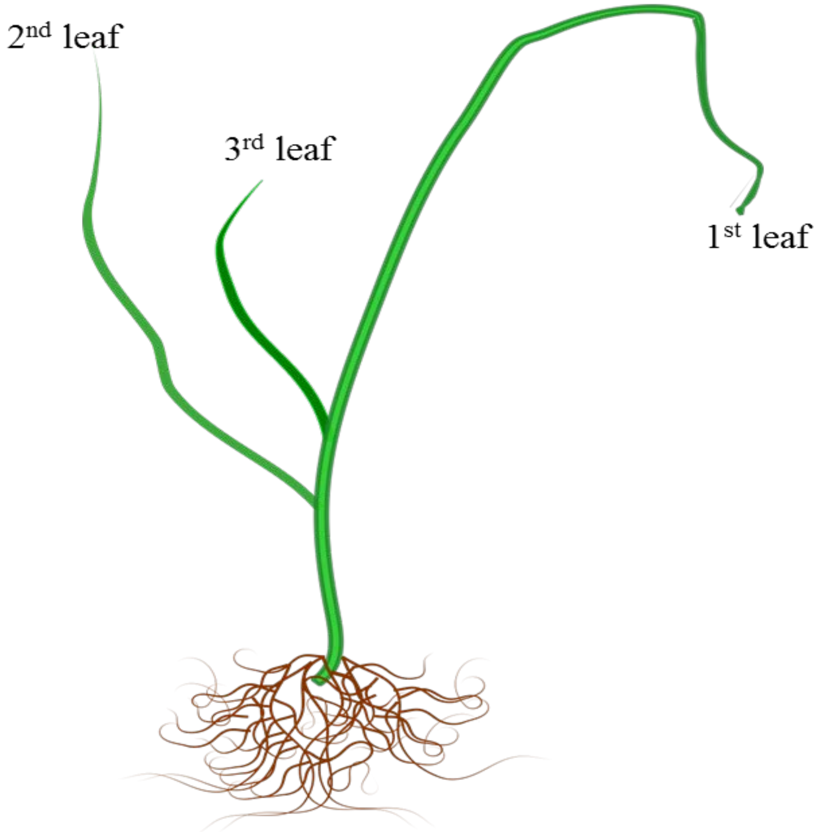


**Figure S1** Cartoon of wheat seedling at 50% of third leaf emergence stage

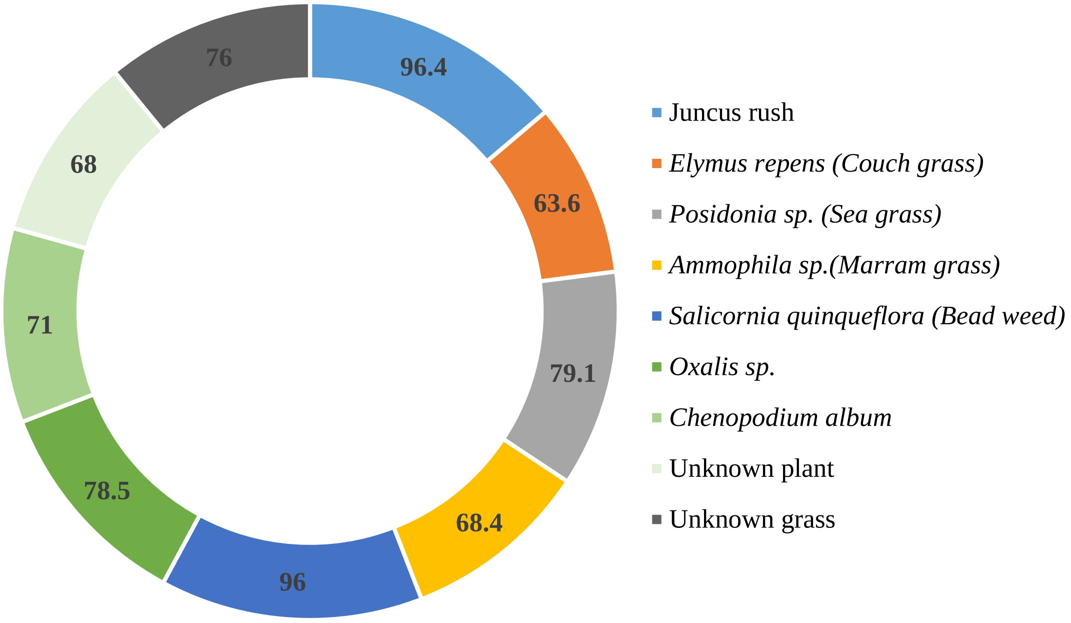


**Figure S2** Colonization frequency of fungal endophytes expressed as percentage (%). The values indicate the percentage of tissue fragments of halophytic species that yielded endophytes in PDA medium.

**Table S2** Fungal isolates from halophytic plants grouped based on their tolerance to 1 M NaCl

| Radial growth inhibition | Reaction type | Number of isolates | Endophytic fungal isolates |
|---|---|---|---|
| >75 % -100 % | Sensitive | 6 | JGE-47, MGE-93, MGE-117, SLE-2, CHE-36, OXE-16 |
| >50 % -75 % | Moderately tolerant | 58 | JGE-48, JGE-50, JGE-50, JGE-51, JGE-53, JGE-54, JGE-59, JGE-109. JGE-110, JGE-112, JGE-114, JGE-115, JGE-116, JGE-118, SGE-45, SGE-63, SGE-100, SGE-134, GE-73, CGE-77, CGE-78, CGE-88, CGE-89, CGE-90, CGE-140, CGE-143, MGE-29, MGE-78, MGE-80, MGE-94, MGE-95, MGE-105, MGE-107, MGE-108, MGE-119, MGE-121, MGE-127, MGE-130, MGE-131, MGE-135, MGE-136, MGE-154, MGE-155, MGE-156, MGE-157, SLE-9, SLE-20, SLE-21, SLE-26, OXE-11, OXE-12, CHE-38, E-68, E-96, E-98, E-125, E-129, E-137, E-138 |
| >25 % - 50 % | Tolerant | 39 | JGE-49, SGE-42, SGE-43, SGE-44, SGE-46, SGE-102, SGE-103, SGE-104, CGE-82, CGE-83, CGE-84, CGE-92, CGE-132, CGE-141, CGE-142, CGE-144, MGE-30, MGE-31, MGE-32, MGE-79, MGE-126, MGE-147, MGE-148, MGE-153, SLE-3, SLE-4, SLE-5, SLE-7, SLE-8, SLE-22, OXE-14, OXE-15, OXE-17, OXE-24, CHE-1, CHE-34, E-65, E-72, E-97 |
| 0-25 % | Highly tolerant | 27 | JGE-52, JGE-55, JGE-58, SGE-60, SGE-61, SGE-62, SGE-85, SGE-86, SGE-87, SGE-101, SGE-133, CGE-74, CGE-75, MGE-81, MGE-106, SLE-6, SLE-10, SLE-19, SLE-27, SLE-28, OXE-13, OXE-25, OXE-37, OXE-40, CHE-39, E-124, E-139 |

**Table S3.** Endophytic fungal isolates identified in the current study based on their ITS sequences, their source of isolation and corresponding GenBank accessions

| S. No. | Isolate code | Host species | Tissue origin | Predicted identity | Query cover (%) | Identity (%) | GenBank Accession number of ITS region |
|---|---|---|---|---|---|---|---|
| 1. | JGE-55 | *Juncus acutus* | Root | *Phomopsis columnaris* | 99 | 99 | MK431041 |
| 2. | JGE-113 | *J. acutus* | Root | *Clonostachys rosea* | 100 | 99 | MK431042 |
| 3. | JGE-115 | *J. acutus* | Root | *Penicillium canescens* | 95 | 99 | MK431043 |
| 4. | JGE-116 | *J. acutus* | Root | *Fusarium sambucinum* | 96 | 98 | MK431044 |
| 5. | JGE-118 | *J. acutus* | Root | *Penicillium canescens* | 96 | 96 | MK431045 |
| 6. | SGE-60 | *Posidonia australis* | Rhizome | *Aspergillus ochraceus* | 100 | 99 | MK431046 |
| 7. | SGE-61 | *P. australis* | Rhizome | *Aquanectria penicillioides* | 95 | 97 | MK431047 |
| 8. | SGE-62 | *P. australis* | Rhizome | *Phaeosphaeria culmorum* | 100 | 99 | MK431048 |
| 9. | SGE-85 | *P. australis* | Rhizome | *Fusarium anthophilum* | 100 | 99 | MK431049 |
| 10. | SGE-86 | *P. australis* | Rhizome | *Plectosphaerella cucumerina* | 99 | 99 | MK431050 |
| 11. | SGE-101 | *P. australis* | Rhizome | *Paraconiothyrium cyclothyrioides* | 100 | 95 | MK431051 |
| 12. | SGE-103 | *P. australis* | Rhizome | *Penicillium brevicompactum* | 94 | 99 | MK431052 |
| 13. | SGE-104 | *P. australis* | Rhizome | *Fusarium anthophilum* | 100 | 99 | MK431053 |
| 14. | SGE-133 | *P. australis* | Rhizome | *Fusarium lacertarum* | 100 | 98 | MK431054 |
| 15. | CGE-74 | *Elymus repens* | Root | *Chaetomium globosum* | 96 | 100 | MK431055 |
| 16. | CGE-75 | *E. repens* | Root | *Paraphaeosphaeria* sp. | 93 | 99 | MK431056 |
| 17. | CGE-83 | *E. repens* | Root | *Fusarium sambucinum* | 100 | 98 | MK431057 |
| 18. | CGE-142 | *E. repens* | Root | *Trichoderma atroviride* | 96 | 95 | MK431058 |
| 19. | MGE-30 | *Ammophila arenaria* | Stolon | Unknown Fungal sp. | 100 | 96 | MK431059 |
| 20. | MGE-80 | *A. arenaria* | Stolon | *Didymella macrostoma* | 100 | 99 | MK431060 |
| 21. | MGE-81 | *A. arenaria* | Stolon | *Alternaria infectoria* | 96 | 98 | MK431061 |
| 22. | MGE-106 | *A. arenaria* | Stolon | Unknown fungal sp. | 99 | 93 | MK431062 |
| 23. | MGE-107 | *A. arenaria* | Stolon | *Microsphaeropsis arundinis* | 100 | 99 | MK431063 |
| 24. | MGE-126 | *A. arenaria* | Stolon | Unknown fungal sp. | 100 | 88 | MK431064 |
| 25. | MGE-148 | *A. arenaria* | Stolon | *Didymosphaeria variabile* | 100 | 96 | MK431065 |
| 26. | MGE-153 | *A. arenaria* | Stolon | Unknown Fungal sp. | 99 | 92 | MK431066 |
| 27. | SLE-3 | *Salicornia quinqueflora* | Root | *Fusarium chlamydosporum* | 100 | 98 | MK431067 |
| 28. | SLE-4 | *S. quinqueflora* | Root | *Fusarium equiseti* | 100 | 98 | MK431068 |
| 29. | SLE-6 | *S. quinqueflora* | Root | *Alternaria chlamydospora* | 100 | 100 | MK431069 |
| 30. | SLE-7 | *S. quinqueflora* | Root | Unknown fungal sp. | 100 | 93 | MK431070 |
| 31. | SLE-8 | *S. quinqueflora* | Root | *Alternaria chlamydospora* | 100 | 100 | MK431071 |
| 32. | SLE-10 | *S. quinqueflora* | Root | *Chaetomium globosum* | 96 | 99 | MK431072 |

| | | | | | | | |
|---|---|---|---|---|---|---|---|
| 33. | SLE-19 | *S. quinqueflora* | Root | *Microsphaeropsis arundinis* | 100 | 99 | MK431073 |
| 34. | SLE-20 | *S. quinqueflora* | Root | *Alternaria molesta* | 100 | 99 | MK431074 |
| 35. | SLE-26 | *S. quinqueflora* | Root | *Alternaria chlamydospora* | 100 | 99 | MK431075 |
| 36. | SLE-28 | *S. quinqueflora* | Root | *Fusarium equiseti* | 100 | 95 | MK431076 |
| 37. | OXE-11 | *Oxalis pes-caprae* | Root | *Fusarium verticillioides* | 96 | 100 | MK431077 |
| 38. | OXE-12 | *O. pes-caprae* | Root | *Alternaria chlamydospora* | 100 | 100 | MK431078 |
| 39. | OXE-14 | *O. pes-caprae* | Root | *Chaetomium globosum* | 100 | 99 | MK431079 |
| 40. | OXE-15 | *O. pes-caprae* | Root | *Fusarium longipes* | 100 | 99 | MK431080 |
| 41. | OXE-17 | *O. pes-caprae* | Root | *Chaetomium globosum* | 99 | 99 | MK431081 |
| 42. | OXE-24 | *O. pes-caprae* | Root | *Fusarium longipes* | 100 | 98 | MK431082 |
| 43. | OXE-37 | *O. pes-caprae* | Root | *Penicillium attenuatum* | 100 | 99 | MK431083 |
| 44. | OXE-40 | *O. pes-caprae* | Root | *Plectosphaerella plurivora* | 100 | 98 | MK431084 |
| 45. | CHE-1 | *Chenopodium album* | Root | *Sordaria tamaensis* | 100 | 98 | MK431085 |
| 46. | CHE-34 | *C. album* | Root | *Penicillium janczewskii* | 96 | 99 | MK431086 |
| 47. | CHE-36 | *C. album* | Root | *Microascus cirrosus* | 96 | 98 | MK431087 |
| 48. | CHE-38 | *C. album* | Root | *Alternaria limaciformis* | 100 | 99 | MK431088 |
| 49. | CHE-39 | *C. album* | Root | *Penicillium* sp. | 100 | 98 | MK431089 |
| 50. | E-65 | Unidentified grass | Stolon | *Alternaria* sp. | 100 | 98 | MK431090 |
| 51. | E-67 | Unidentified grass | Stolon | *Setosphaeria rostrata* | 99 | 100 | MK431091 |
| 52. | E-97 | Unidentified grass | Stolon | *Bipolaris sorokiniana* | 100 | 99 | MK431092 |
| 53. | E-124 | Unidentified brassica | Root | Unknown fungal sp. | 48 | 92 | MK431093 |
| 54. | E-139 | Unidentified brassica | Root | *Fusarium sterilihyphosum* | 96 | 98 | MK431094 |

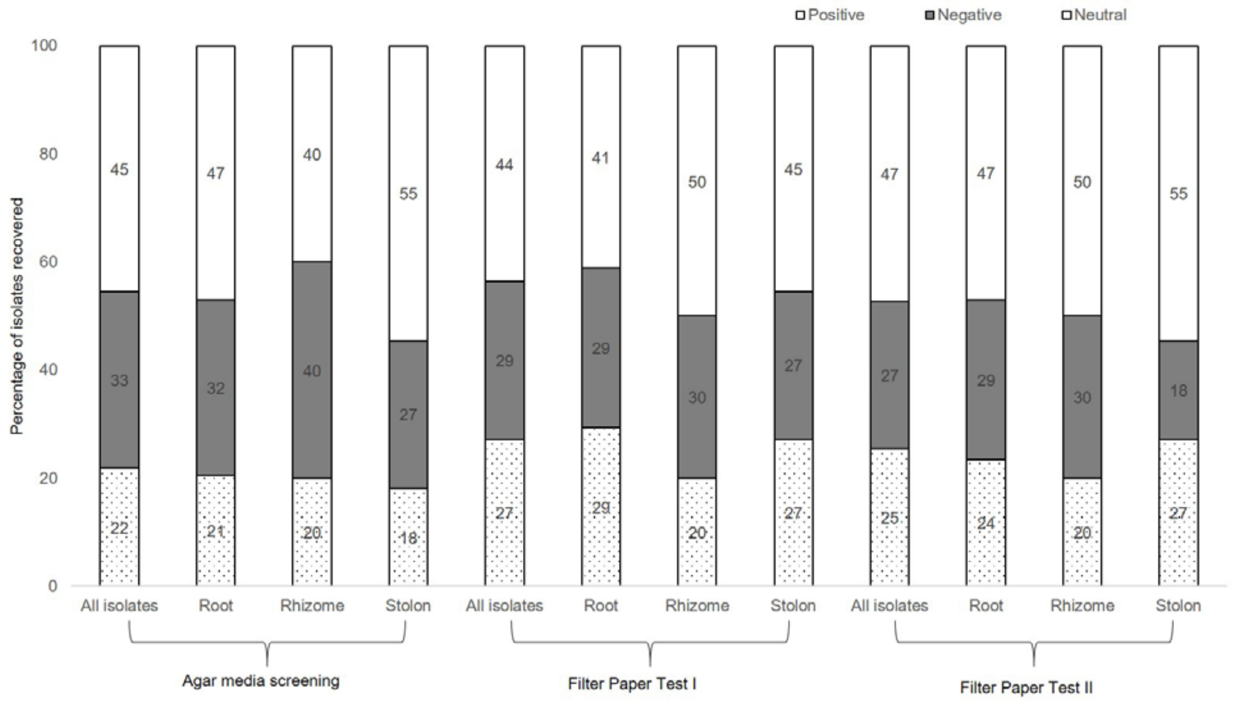


**Figure S3** Percentage of endophytic isolates recovered from different tissues showing differential reactions when co-cultivated with wheat seedlings and subjected to **(a)** agar media screening, **(b)** filter paper I and **(c)** filter paper II screening, to study the moisture deprivation tolerance. The number inside each stack bar indicates the relative percentage of endophyte isolates corresponding to particular tissue with their reaction **(Positive**= enhanced tolerance, **Negative** = decreased tolerance or retarded growth upon inoculation, **Neutral** = no measurable influence)

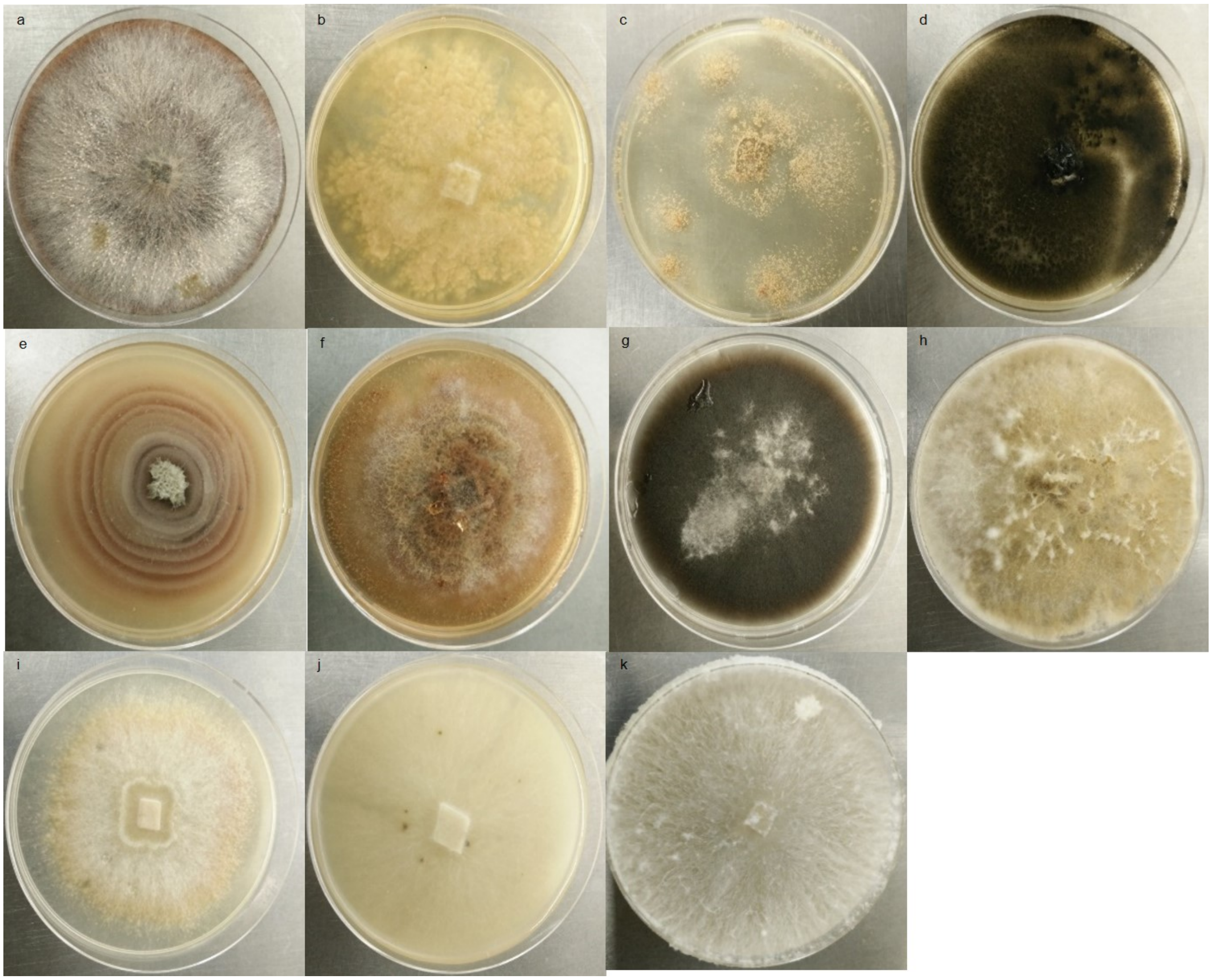


**Figure S4** Pure cultures of fungal endophytic isolates used for salt tolerance studies on wheat seedlings a) Unknown fungal sp. (MGE-106) b) *Chaetomium globosum* c) *Aspergillus ochraceus* d) *Alternaria chlamydospora* e) *Microsphaeropsis arundinis* f) *Chaetomium globosum* g) *Alternaria infectoria* h) *Didymosphaeria variabile* i) *Aquanectria penicillioides* j) *Chaetomium globosum* k) *Trichoderma atroviride*
